# Development of an integrated genomic map for a threatened Caribbean coral (*Orbicella faveolata*)

**DOI:** 10.1101/183467

**Authors:** Jacob Snelling, Katherine Dziedzic, Sarah Guermond, Eli Meyer

## Abstract

Genomic methods are powerful tools for studying evolutionary responses to selection, but the application of these tools in non-model systems threatened by climate change has been limited by the availability of genomic resources in those systems. High-throughput DNA sequencing has enabled development of genome and transcriptome assemblies in non-model systems including reef-building corals, but the fragmented nature of early draft assemblies often obscures the relative positions of genes and genetic markers, and limits the functional interpretation of genomic studies in these systems. To address this limitation and improve genomic resources for the study of adaptation to ocean warming in corals, we’ve developed a genetic linkage map for the mountainous star coral, *Orbicella faveolata*. We analyzed genetic linkage among multilocus SNP genotypes to infer the relative positions of markers, transcripts, and genomic scaffolds in an integrated genomic map. To illustrate the utility of this resource, we tested for genetic associations with bleaching responses and fluorescence phenotypes, and estimated genome-wide patterns of population differentiation. Mapping the significant markers identified from these analyses in the integrated genomic resource identified hundreds of genes linked to significant markers, highlighting the utility of this resource for genomic studies of corals. The functional interpretations drawn from genomic studies are often limited by the availability of genomic resources linking genes to genetic markers. The resource developed in this study provides a framework for comparing genetic studies of *O. faveolata* across genotyping methods or references, and illustrates an approach for integrating genomic resources that may be broadly useful in other non-model systems.

## INTRODUCTION

Genomic approaches enabled by high-throughput DNA sequencing technologies are powerful tools for studying the genomic basis of complex traits and adaptive responses to selection [1,2]. Once restricted to the few model systems where large research communities supported development of extensive genomic resources, advances in DNA sequencing technologies have recently brought the same approaches within the reach of researchers studying ecological or evolutionary questions in non-model systems [3-6]. The ability to simultaneously genotype large numbers of genetic markers without prior sequence information [7,8] allows researchers to conduct genome-wide association studies and QTL mapping in non-model species, identifying genomic regions associated with the trait of interest.

It’s become relatively easy to generate large genomic datasets from any species, but interpreting these data has remained challenging, in the non-model species of interest for many ecological and evolutionary questions, because of the limited genomic resources available in these systems. For example, reef-building corals are severely threatened by rising ocean temperatures [9,10], and the development of genomic resources in these systems has historically lagged behind better studied systems like model or crop species. Transcriptome assemblies are now available for many corals [11-17], but while these collections of gene sequences are useful for many studies they offer no information on their positions in the genome. Draft genome assemblies are available for a smaller number of systems [18-20], but in most cases these assemblies have remained fragmented (thousands of scaffolds), lacking the long-range structural information provided by chromosome-scale assemblies. The limited structural information in these resources obscures relationships between genetic markers and genes, leaving considerable uncertainty for functional interpretation of genomic studies in corals.

Linkage mapping provides structural information that complements the limitations of sequence assemblies. In this approach, the relative positions of genetic markers in the genome are inferred from their co-segregation in a mapping population (typically a collection of full-sibling individuals with known parental and gandparental genotypes). This approach was developed in model systems [21] and has been widely used in those systems and a wide range of crop species to study genomic basis for trait variation [22]. In non-model species studied for ecological and evolutionary questions, linkage analysis has been limited by the difficulties of conducting controlled genetic crosses and the scarcity of genetic markers in these systems. In experimentally tractable model or crop species, linkage analysis has historically relied on advanced multi-generation pedigrees or inbred lines that can’t be realistically achieved in slow-growing or difficult to cultivate species. However, researchers have developed methods for linkage analysis in first-generation crosses between parents sampled from natural populations [23,24]. Several methods are now available for genotyping large numbers of genetic markers (single nucleotide polymorphisms, SNPs) without prior sequencing information or marker development [7,8,25,26]. Together, these approaches have enabled development of genetic linkage maps for several non-model systems studied for ecological or evolutionary questions [27-30], including our previous studies of an Indo-Pacific coral (*Acropora millepora*) [31,32].

Scleractinian corals (Cnidaria) form the physical and ecological foundation of coral reefs, supporting some of the most diverse and productive of all marine ecosystems [33,34]. Unfortunately, reef-building coral populations have declined globally [35-37] and their persistence is threatened by ongoing climate change. These corals live in obligate associations with symbiotic dinoflagellates (Symbiodinium sp.), and this association is highly sensitive to warming, breaking down during thermal stress in the well-known phenomenon of coral bleaching [38,39]. The increasing degradation of global coral populations, resulting at least in part from ocean warming, has led to grave concerns over the fate of corals and the reef ecosystems they support during ongoing climate change [9,10,40].

The question of whether (and how quickly) corals can adapt to ocean warming has risen to the forefront of coral biology [41,42]. Predicting these adaptive responses requires an understanding of existing genetic variation in the traits under selection [22]. To study this variation, biologists are increasingly turning to genomic approaches in many non-model systems including corals [43,44]. Genome sequence assemblies are available or underway for corals for several species [18,20]. However, genetic linkage maps, which provide long-range structural information required to identify regions under selection, have only been described for Indo-Pacific species [26,31]. The Caribbean has a unique and highly degraded coral fauna [36], and linkage maps remain unavailable for any Caribbean species, limiting the scope of genome studies that can be conducted in these systems. Studies of adaptation are based on genetic variation, which is population specific and cannot be reliably generalized across populations within a species, let alone generalized across the deep divergence (>24 mya) between Caribbean and Indo-Pacific species [45]. Understanding adaptive potential in Caribbean corals requires improvement of genomic resources for these species and populations.

To enable these studies, we’ve developed a genetic linkage map using larvae from controlled crosses of the important Caribbean reef-builder *Orbicella faveolata*. To illustrate the usefulness of this resource for genomic studies in *O. faveolata*, we’ve conducted a demonstration study using a collection of corals housed in our research aquarium to study trait associations and genetic differentiation. We’ve integrated the linkage map with publicly available genome and transcriptome assemblies to provide a common frame of reference for genomic studies in this widely studied and ecologically important component of Caribbean reefs.

## MATERIALS AND METHODS

### Genetic linkage analysis

#### Genetic crosses

To develop a mapping population for linkage analysis, we conducted controlled genetic crosses between parental corals sampled from a natural population.

For these crosses, we used gametes from four parental colonies collected prior to the annual mass-spawning event in 2012 (Flower Garden Banks, Gulf of Mexico, Permit number FGBNMS-2012-002). On the night of the natural spawning event, we conducted experimental crosses as we’ve previously described for other coral species [46,47] and as others have described for *Orbicella* [48,49]. We collected sperm and eggs from colonies isolated prior to the natural spawning event, and combined these in a factorial design to produce six different crosses. After spawning and fertilization, we sampled tissue from each parental colony (preserved in 100% ethanol) for later analysis of parental genotypes.

To maintain larval cultures at appropriate temperatures on board the R/V Manta (Galveston, TX), we submerged culture vessels in flow-through water baths constantly exchanged with ambient seawater. We sampled pools of >200 larvae at 24 hours postfertilization, late enough in development to allow multiple rounds of cell division and ensure sufficient DNA content for genetic analysis, but early enough to minimize segregation distortions from allele-specific larval mortality. Each of these samples could in principle be used as a mapping population for linkage analysis.

#### SNP genotyping parental colonies

To choose the most informative cross for genetic linkage analysis, we conducted multilocus SNP genotyping of parental corals to identify the most genetically dissimilar pair. For this purpose we used a sequencing based approach (2bRAD) [26] we’ve previously used in other coral species [31,50]. This is one of a growing family of methods that sequence a defined subset of the genome to simultaneously discover and genotype thousands of SNP markers [7,8,25]. RAD refers to the selection of genomic regions using restriction endonucleases (restriction associated DNA, RAD) [25], and “2b” refers to the type IIb enzymes used for library preparation [26]. In this approach, libraries can be easily customized to target a defined subset of restriction sites using selective adaptors. To profile genetic variation among parents, we prepared sequencing libraries as described in [26], using *Alf*I restriction endonuclease for all libraries in this study. We combined libraries in equimolar amounts for multiplex sequencing on HiSeq 2000 platform at Oregon State University’s Center for Genome Research and Biocomputing.

#### SNP genotyping individual larvae

To study genetic linkage among SNP markers in this population, we conducted multilocus SNP genotyping of individual larvae from the chosen family. First, we extracted DNA from individual larvae (preserved in 100% ethanol) using a custom procedure designed to maximize yield from these small larvae (<200 μm in length). We first lysed individual larvae in 10 μl of a lysis buffer prepared from Buffer TL (Omega Bio-tek, Product No. PD061) with OB Protease (Omega Bio-tek, Product No. AC114) and RNAse A (Qiagen) each at 2 μg μl^-1^. We incubated samples in lysis buffer for 15 minutes at 55°C to fully lyse larval tissues, then precipitated nucleic acid from the lysates with isopropanol, adding 5 μg glycogen as a co-precipitant to improve yields. To minimize co-precipitation of contaminants from coral tissues that can inhibit enzymatic reactions, we conducted these precipitations with minimal alcohol concentrations (1 volume isopropanol) and incubated at room temperature rather than at freezing temperatures. Finally, we dissolved pellets after the first precipitation in 10 mM Tris (pH 8.0) and repeated the precipitation to further purify the DNA and minimize enzymatic inhibitors.

We prepared 2bRAD genotyping libraries from 128 individual larvae, focusing on the cross between the two most genetically dissimilar parents. 110 of these samples produced successful sequencing libraries, which we quantified in equimolar ratios based on qPCR quantification of each library. We quantified these libraries using Sensifast SYBR master mix (Bioline; Boston, MA), and the PCR primer sequences shown in the Illumina Customer Sequence Letter (Lib-1: AATGATACGGCGACCACCGA and Lib-2: CAAGCAGAAGACGGCATACGA). To minimize sequencing costs, we prepared reduced representation libraries using selective adaptors designed to capture 1/4 of AlfI restriction sites in these libraries (adaptors ending with “NR“ overhangs) [26]. We combined libraries in equimolar amounts for multiplex sequencing on two lanes of the HiSeq 2000 platform at Oregon State University’s Center for Genome Research and Biocomputing.

#### Sequence analysis

Before using these sequences to determine genotypes in each sample, we first processed the data to eliminate low-quality or uninformative reads. We processed reads essentially as described in [26,50]. We first truncated reads to remove sequences derived from sequencing adaptors, then excluded low quality reads having ≥ 3 positions with quality scores < 20. We also removed reads matching known adaptor sequences used during library preparation (Smith-Waterman alignment scores > 18). (All scripts used for these processing steps are publicly available at https://github.com/Eli-Meyer/sequence_processing). The high quality sequences that remained after filtering formed the basis for all subsequent analysis.

To determine SNP genotypes in each sample, we followed the same approach we have previously described [26] with minor modifications described here. 2bRAD data can be analyzed using a reference genome or by constructing a reference de novo by clustering reads from the samples of interest [26]. This is conceptually similar to the approach taken in STACKS [51], a software package widely used for other kinds of RAD data, which identifies loci by clustering sequence data to develop a catalog of reference loci, then aligns reads from each sample to this reference to genotype each locus in each sample. To provide an independent comparison with the draft genome assembly, we chose to use this *de novo* approach rather than align reads to the draft assembly. To that end, we developed a reference by first identifying sequences observed at least twice in the highest quality reads (quality scores> 30 at all position), further clustering these sequences to identify groups of related alleles differing by no more than 2 base pairs, and finally using relationships among sequences in each cluster to identify sub-clusters of related alleles differing by no more than 1 base pair. The consensus sequence from each of the final sub-clusters was chosen as the reference for each locus, and these sequences combined to provide a reference for mapping and genotype calling. Scripts automating this process are archived at (https://github.com/Eli-Meyer/2brad_utilities).

Next, we mapped all reads from each sample against this reference, providing a common framework for evaluating genetic variation among samples. We mapped reads using SHRiMP2 [52], rejecting ambiguous alignments, alignments shorter than 34 bp, or alignments with fewer than 32 matching bases (out of 36). We then determined genotypes at each locus with ≥ 10× sequencing coverage, based on nucleotide frequency thresholds [26]. Here, we called loci homozygous if any alternate alleles were present at ≤1% allele frequency, and heterozygous if alternate alleles were present at ≥25% allele frequency. Loci with intermediate allele frequencies were left undetermined, to avoid genotyping errors occurring in loci with these frequencies (sacrificing number of genotypes to minimize genotyping error). This process provided a matrix of high-confidence SNP genotypes, describing sequencing variation across thousands of SNP loci (rows) in multiple samples (columns).

#### Linkage analysis

To understand linkage among these markers, and develop a genetic linkage map reflecting their physical arrangement on chromosomes, we analyzed co-segregation of genotypes in each pair of markers. We conducted all linkage analysis using the R package *Onemap* [23], which was developed to take advantage of multiple marker configurations in F1 crosses between parents collected from outbred natural populations. The core of this method is the simultaneous estimation of linkage and linkage phases using maximum likelihood [24]. Unlike previous pseudo-testcross strategies, this approach handles both partially and full-informative markers and produces a single map rather than parent-specific maps.

Prior to linkage analysis, we filtered the SNP genotypes to minimize missing data, genotyping errors, and markers unsuitable for mapping. We first excluded samples genotyped at too few loci (<400 SNPs; removing 18 of 112 samples) because of low sequencing coverage or library qualities. Next we excluded loci genotyped in fewer than 30 samples, as an absolute minimum number of samples for linkage analysis. We excluded tags containing excessive sequence variation (>2 SNPs per 36-bp tag), since these may reflect errors introduced by repetitive genomic regions. For any tags containing more than one SNP, we selected a single representative SNP for statistical analysis (since such tightly linked SNPs would effectively segregate as a single locus and cannot be considered independent).

Since information on parental genotypes is required for linkage analysis, we further filtered the SNP genotypes to exclude loci genotyped in neither parent. Finally, we excluded samples deviating from expected segregation ratios at too many loci, and markers showing extreme deviations from expected segregation ratios across samples (average deviations in allele frequencies ≥0.3 for both filters). This stringent selection produced a set of high-quality SNP genotypes that served as the basis for all linkage analysis, and satisfied initial tests for valid segregation patterns implemented in the mapping software.

To identify groups of linked markers corresponding to chromosomes, we estimated two-point recombination frequencies for each pair of markers, requiring LOD scores ≥8 and recombination frequencies ≤0.5 to establish linkage. We assigned markers to linkage groups using these estimates and the transitive property of linkage, and required at least 20 markers per linkage group. We then estimated the arrangement of markers within each linkage group using an iterative mapping procedure implemented in the *Onemap* package. First, we established an initial framework for each linkage group by exhaustively comparing all arrangements among 6 markers (LOD threshold ≥ 3). Next, we sequentially attempted to add the remaining markers to this map using a touchdown procedure with an initial round using a LOD ≥ 3 threshold, followed by a subsequent round at LOD ≥ 2. Finally, we added the remaining unmapped markers at the positions best supported by recombination frequencies relative to markers already mapped.

After constructing each linkage group using this automated procedure, we identified misplaced markers by visualizing patterns of LOD scores and recombination frequencies among mapped markers. To eliminate any effects of these problematic markers on the map, we excluded these using the *drop.marker* function and reconstructed the map (as above) without them, then attempted to reintroduce them to the map using the *try.seq* function. If this succeeded in incorporating the problematic markers without distorting map lengths or introducing large gaps, we retained the markers in these new positions, discarding them otherwise. This analysis placed our SNP markers into a series of linkage groups which we interpret as corresponding to chromosomes (or portions of chromosomes) in the *O. faveolata* genome.

### Integrating the linkage map and genome assembly

To develop a framework for genomic analysis of traits and populations in *O. faveolata*, we integrated the linkage map and genome assemblies as shown in Fig 1. For this study, we initially developed an integrated map using a draft assembly that Monica Medina and Bishoy Hannah kindly shared with us prior to publication, and the transcriptome assembly described in [11]. When the updated assembly became available (NCBI accession GCF_002042975.1), we updated the integrated map using that assembly to develop the resource reported here. We compared 2bRAD tags (36-bp *AlfI* fragments) from the *de novo* reference with the genome assembly using the same mapping approaches previously described for analysis of 2bRAD sequencing libraries (above), discarding ambiguous or weak (>1 mismatch) alignments.

**Fig 1.**
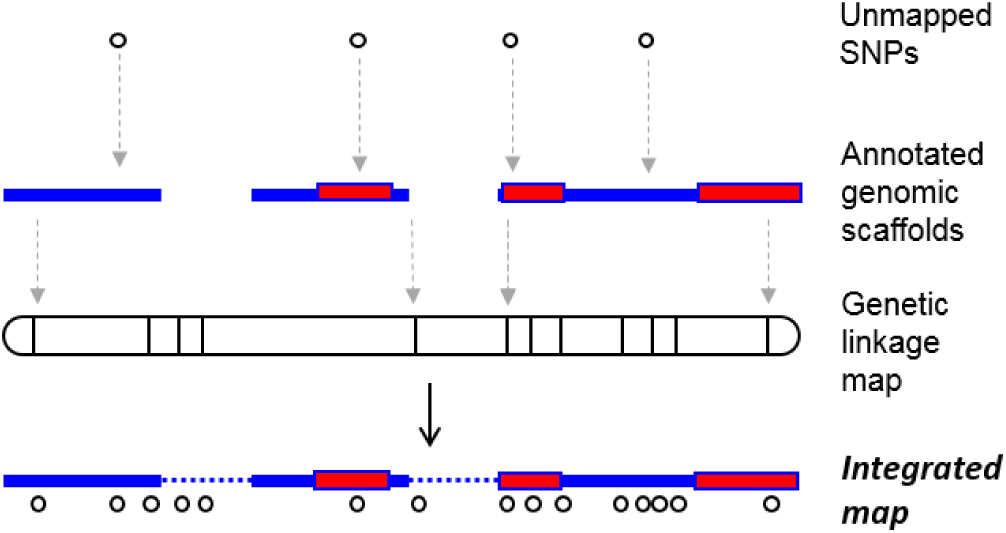
An overview of our approach for combining genetic linkage map and genome assembly to place SNP markers in an integrated map. Genetic markers (RAD tags) from the linkage map are mapped to genomic scaffolds (blue bars) to identify the relative positions of these scaffolds in the genome, then additional SNP markers (open symbols) are mapped to these scaffolds. The integrated map describes the relative positions of genetic markers and genes, uncovering information that wouldn’t be apparent from the constituent datasets alone.

Next, we used the linkage map to infer the relative positions of genomic scaffolds in the genome. We used map positions to order scaffolds in each linkage group, and for scaffolds containing two or more markers, to orient the scaffolds (forward or reverse). Because most scaffolds in this assembly are smaller than the genetic distances that could be resolved with the scale of sampling used here, we chose not to infer genetic distances for each marker within a scaffold, instead assigning each scaffold a single position in the genetic map. To evaluate agreement between the linkage map and genome assembly, we focused on scaffolds containing multiple mapped markers. For each scaffolds containing multiple mapped markers, we asked whether all markers were assigned to a single linkage group, counting these as agreement between the linkage map and assembly. If markers were mapped to more than one linkage group, we assigned the scaffold to the consensus linkage group if there were at least twice as many markers assigned to that group as the second most frequent group. Ties (scaffolds equally strongly associated with more than one linkage group) were discarded. This conservative approach discarded areas of disagreement between the linkage map and assembly rather than attempting to declare one of the resources correct. Finally, we collected additional SNP markers that could not be directly mapped (either because they were not informative for mapping or received inadequate sequencing coverage in the mapping population), and placed these based on their matches to scaffolds included in the integrated map.

This integrated map provides a framework for comparing genomic studies of *O. faveolata* (previous or future) using either the linkage map, the genome assembly, or the previously reported transcriptome assembly [11] as a reference. The initial version developed in this study is not a final product but a starting point, and we anticipate that further developments in any of the constituent databases will feed back to further improve the integrated map. We describe a standardized procedure for integrating these resources, and present an initial version of the integrated map based on current resources. Our vision is that this can be readily updated as the resources develop, and the updated versions made publicly available for use by the coral research community.

### Genomic studies enabled by genetic linkage map

To illustrate the kind of analyses enabled by this integrated genomic resource, we conducted several small-scale demonstration studies using a collection of coral specimens available at the time in our research aquariums. The scale and sampling design of these studies was not intended to support broad biological conclusions about natural populations, but to illustrate the utility of this resource for genomic studies in *O. faveolata*. To that end, we measured variation in bleaching responses during a controlled thermal stress experiment, then tested for associations between SNP genotypes and bleaching responses. Because these corals presented conspicuous color variation, we also tested for genetic associations with variation in color (fluorescence from green fluorescent proteins naturally produced by the coral host) [53,54]. To identify genomic regions differentiated among populations, we compared estimates of genetic differentiation at each locus with the overall genomic background.

The structural information gained from the linkage map benefits these studies in two ways. (i) Considering the relative positions of these markers determines whether SNPs statistically associated with the trait or population of interest are physically linked in one or a few regions, or scatted across the genome. (ii) Considering the relative positions of expressed sequences and genetic markers provides information on the genes linked with significant SNPs identified in each study, which may provide insights into functional differences between the groups being compared.

#### Genomic regions associated with bleaching responses

To identify genomic regions associated with bleaching responses, we tested for effects of genotype on variation in bleaching responses during a controlled thermal stress experiment.

#### Preparation and sampling of corals

To study genomic patterns associated with traits and population differentiation, we prepared an array of fragments from a collection of coral genotypes (*O. faveolata*) available at the time in our research aquariums. These included 12 fragments sampled from colonies in Florida Keys (obtained by our colleague Rebecca Vega-Thurber under permit number FKNMS-2013-016 and kindly shared with us for this study) and 12 fragments collected in 2013 from the Flower Garden Banks National Marine Sanctuary (collected under permit FGBNMS-2013-004 to Meyer). We prepared replicate fragments from each colony for biological replication of bleaching responses in each genotype, and maintained these fragments in research aquaria for >2 months for recovery and growth prior to thermal stress experiments. During this period we maintained corals in a large recirculating research aquarium (total system volume > 2500 L) at 27°C, under approximately 150 μmol photons m^-2^ s^-1^ with salinity maintained at 35 ppt and pH maintained between 8.0 – 8.2. To measure bleaching responses, we randomly distributed five fragments from each colony into each of two recirculating aquarium systems (total volume approximately 250 liters each) held initially at 27°C. We then imposed a warming treatment in one room, increasing temperature gradually by 0.5°C per day to a maximum of 32°, then maintaining that temperature, while the other room was maintained at a constant 27°C.

We monitored bleaching progress by daily visual inspection, terminating the experiment and sampling when bleaching was visible in approximately half the fragments in the thermal stress treatment. We documented bleaching responses by photographing each fragment in constant conditions under light microscopy. For comparison with field studies, we quantified bleaching responses using the same approach widely used in field surveys [55]: comparison with a standardized color card [56]. To estimate bleaching responses in each colony, we assigned each fragment a color score in this way and calculated bleaching responses as the difference between average scores of stressed and control fragments from each colony.

*Accounting for variation in symbiont types.* Because variation in the thermal tolerance of the coral holobiont (the coral plus its algal and microbial symbionts) [57-59] can be influenced by the symbiont community [60-62], we investigated the symbiont community in each sample to account for this confounding factor. We measured the dominant symbiont type in each sample using Sanger amplicon sequencing. For this purpose we used primers ITS2F (GAATTGCAGAACTCCGTG) and ITS2R (GGATCCATATGCTTAAGTTCAGCGGGT) to amplify a region in the internal transcribed spacer of the ribosomal gene array previously described for analysis of diversity in algal symbionts of Cnidarians [63]. We conducted PCR using PerfectTaq (5 Prime, DNA polymerase according to the manufacturer’s instructions, with both primers at 0.4 μM final concentration, 50 ng DNA template per reaction, and annealing temperatures of 48°C. We removed PCR primers by treating PCR products with 0.5 U Antarctic Phosphatase and 0.6 U Exonuclease I (NEB) at 37°C, then inactivating the enzymes at 80°C and analyzing the resulting amplicons by Sanger sequencing with the ITS2F primer at OSU’s Center for Genome Research and Biocomputing. We classified the dominant symbiont type in each colony by comparing sequences from each sample with representative ITS2 sequences from Symbiodinium clades A-D (NCBI accessions AF333505, AF333506, AF333507, AF333511, AF333512, AF333514, AF334660, JX415807, EU333743, and EU333731). This analysis provided a coarse survey of symbiont communities to account for possible confounding effects of variation in symbiont types on variation in bleaching responses of the coral holobiont. We included this information in statistical models (below) used to test for associations between genetic variation and bleaching responses.

*Statistical analysis.* We partitioned bleaching responses using a linear mixed model to evaluate the relative contributions of geographical origin, symbiont type, and colony to this variation. To test for associations between bleaching responses and genotypes at each locus, we used a mixed model described by [64] and implemented in the R package *rrBLUP* [65]. For this analysis we calculated the additive relationship matrix among samples based on SNP genotypes, and used this matrix as described [66] to control for population structure, which can lead to spurious trait-genotype associations. We used the *GWAS* function in *rrBLUP* to test for associations between bleaching responses and SNP genotypes at each locus. To control for errors arising from conducting multiple tests, we controlled false discovery rates (FDR) at 0.05 as described [67].

#### Genomic regions associated with color variation

The corals used for this study presented conspicuous variation in visible colors resulted from natural variation in the expression or properties of green fluorescent protein (GFP) in these samples. To quantify this variation, we used fluorescence microscopy and image analysis, photographing each coral fragment from the control treatments under fluorescence microscopy at the endpoint of the thermal stress experiment (excitation 470 nm, emission 500-550 nm). We used ImageJ software to quantify fluorescence in these images, as the average intensity in the green channel for areas of each image containing coral tissue after background removal.

*Statistical analysis.* To identify genetic markers associated with fluorescence, we conducted a mixed model analysis at each SNP locus, accounting for genetic relationships among samples as previously described for analysis of bleaching responses. This analysis identified a set of SNPs statistically associated with natural variation in corals’ color (fluorescence) in our test population.

#### Genomic patterns of population divergence

To illustrate the utility of our linkage map for studying genomic patterns of differentiation, we analyzed these patterns in the highly differentiated samples in our collection originating from Texas (Flower Garden Banks, Gulf of Mexico) and Florida (Florida Keys). The large number of loci genotyped in sequencing-based approaches like 2bRAD make it possible to identify strongly-differentiated regions that may result from differences in selection or demographic histories. To that end, we estimated F_S__T_ at each locus and genomewide using the R package diveRsity [68]. For this analysis, we used a sliding window approach (window size = 25 markers) to investigate genomic patterns of genetic differentiation between populations. This analysis allowed us to evaluate whether each region of the genome was unusually differentiated or conserved relative to the overall genomic background.

#### Data availability

We’ve archived data from each stage of our analysis in publicly available records. We archived processed (high-quality) DNA sequences used for genotyping of parental corals, our larval mapping population, and the colonies used for association studies at NCBI’s Sequence Read Archive under accessions SUB2925599, SUB2923030, and SUB2925759 respectively. We’ve provided access to the integrated genomic resource (integrated map) designed here in several locations. To establish a stable record of the version described in this publication, we provide a flat file in Supplementary Information (Supplementary Table S1), and will also maintain updated versions on the author’s laboratory website hosted at Oregon State University (http://people.oregonstate.edu/∼meyere/data.html), updating the current release as the constituent maps and assemblies are updated.

## RESULTS

### Genetic diversity among parental corals

To maximize genetic diversity in our mapping population, we conducted and sampled multiple crosses using gametes from four colonies, then later compared multilocus genotypes of parental corals to identify the combination of parents with the largest number of informative markers for linkage analysis.

To that end, we conducted multilocus SNP genotyping of parental samples using 2bRAD, sequencing 28.4 million reads per sample on average, 81% of which survived filtering for quality and information content (Supplementary Table S2). We developed a de novo reference from these reads as previously described [26], and were able to map nearly all HQ reads (92%) back to this reference. Analysis of these data allowed us to genotype 3 million base pairs in one or more samples. After extensive filtering to minimize missing data and genotyping errors, we identified a set of 14,266 high-quality SNPs in these data, and used these for genetic comparisons among potential parents.

We estimated genetic distance among genotypes based on these SNPs as the proportion of allelic differences in each pairwise comparison. We found genetic distances among corals ranging from 0.19 to 0.38, and chose the most genetically dissimilar pair of corals (C and F), using progeny from this cross for linkage analysis.

### Multilocus SNP genotypes from individual larvae

To enable genomic studies of trait associations or population differentiation in *O. faveolata*, we developed a genetic linkage map using SNP genotypes from this mapping population. To that end, we prepared 2bRAD genotyping libraries from individual larvae. The small size of these samples (approximately 100 μm) introduced challenges for library preparation. We attempted libraries from 128 individuals, succeeding for 110 samples. We recovered sufficient sequencing depth for 92 of these (2.13 million reads per sample on average), genotyping >700 kb across these samples at ≥10x coverage, including 4,322 SNPs (Supplementary Table S2). We developed a new reference from these larval sequences, since these early stages naturally lack algal symbionts that may contribute to sequence data from adult tissues. To compare parental and offspring genotypes for linkage analysis, we reanalyzed the parental data (colonies C and F) using this same reference, and combined all parental and offspring genotypes into a matrix of SNP genotypes. After extensive filtering to minimize missing data and genotyping errors, we identified a set of 2,651 SNPs called with high confidence in most samples and focused on these markers for linkage analysis.

Using these markers, we examined pairwise genetic distances among all samples to screen for contamination during fertilization or larval culture. Nearly all samples (88) matched the expected pattern of half the parental distance (parental distance = 0.48; average expected full-sibling distance = 0.24; observed distances 0.20-0.28). A few samples showed unexpectedly large genetic distances (range: 0.29-0.31), and we excluded these as likely contaminants prior to linkage analysis. After further filtering to exclude SNPs that were not genotyped in either parent, SNPs with uninformative configurations (e.g. AA x BB), and samples or loci showing extreme deviations from expected segregation ratios, we were left with a dataset including 84 individuals and 1,430 high-quality SNPs suitable for linkage analysis.

Our analysis of segregation ratios revealed extensive deviations from expected Mendelian ratios. Filtering for segregation ratios removed a substantial number of otherwise high-quality SNPs (735) because of extreme deviations from expected ratios (difference between expected and observed frequencies > 0.3). After filtering for extreme deviations from expected segregation ratios, the 1,430 high quality SNPs that remained still included markers (432 SNPs) that could not be assigned to linkage groups with high confidence, likely resulting from more subtle distortions in segregation ratios. For the map presented here, we used a stringently filtered subset of 998 SNP markers, choosing to sacrifice marker density for confidence in the overall map.

### Development of a genetic linkage map

We analyzed recombination frequencies among SNP markers to identify a set of linkage groups corresponding to chromosomes, and mapped the relative positions of markers on each group. Of the 1,430 HQ SNPs used for linkage analysis, we were able to assign most (87%) to linkage groups with high confidence (LOD scores ≥ 8). Most of the mapped markers (998, or 81% of mapped markers) were captured in the 16 largest linkage groups (LGs), a biologically plausible number comparable to the range of haploid chromosome numbers (n) reported in other Scleractinian corals (n = 14-27; [69-71]). Because chromosome numbers have not been directly determined in *O. faveolata*, we included all linkage groups containing sufficient markers to provide meaningful information on large-scale genomic structure (>20 markers) rather than selecting a number of groups based on prior expectations. The small groups discarded at this threshold provided little structural information (85% of the discarded groups had <5 markers). Since the number of linkage groups identified from linkage analysis of multilocus SNP genotypes is influenced by the thresholds chosen, we do not interpret the number of linkage groups shown here as a precise estimate of chromosome numbers, aiming instead to capture as much useful structural information as possible from the existing data.

Next, we determined the order of and distance between markers on each linkage group based on recombination frequencies, producing a map with a total length of 2,049 cM, and linkage groups ranging from 47 to 223 cM in length (Fig 2). The average marker interval was 2.05 cM. A majority of markers in this map (67%) were closely linked other markers (≤ 2 cM to nearest neighbor in the map), and most were at least moderately closely linked to another marker (86% of markers within 5 cM of their nearest neighbor). This linkage map provides a framework for organizing genomic sequencing resources, and for high-resolution genomic analysis of trait associations and population divergence in *O. faveolata*.

**Fig 2.**
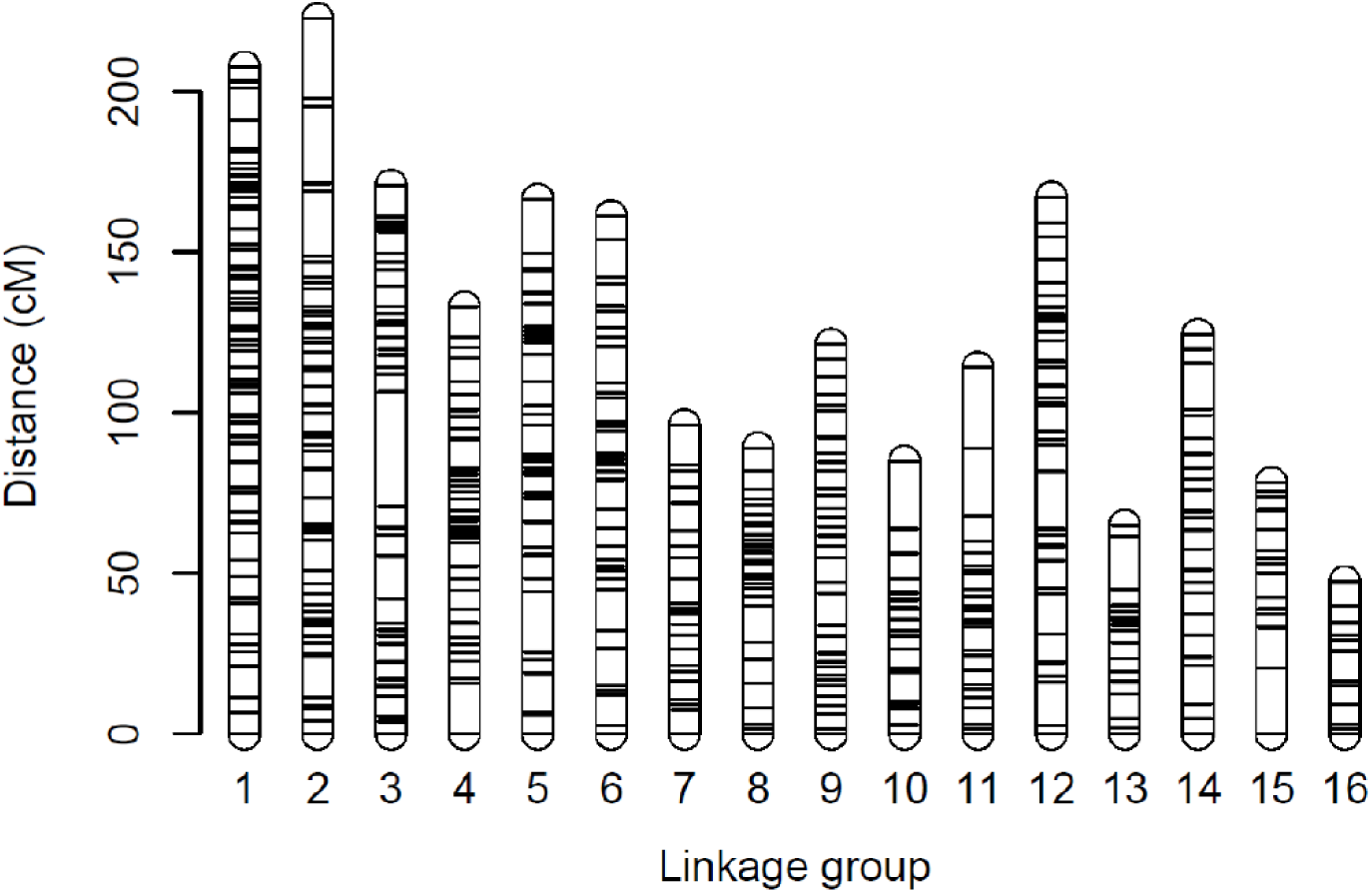
A genetic linkage map based on SNP genotyping of individual larvae from the mountainous star coral *Orbicella faveolata*. This resource organizes 998 genetic markers into 16 linkage groups with a total map length of 2048.5 cM (1 marker per 2.05 cM on average).

### An integrated genomic map for *O. faveolata*

To maximize the utility of these resources for asking biological questions about differences among phenotypes or populations, we combined our genetic linkage map with an annotated genome assembly to develop an integrated map describing the relative positions of genetic markers and genes in the genome of *O. faveolata*. This integrated resource provides a shared framework for comparing genomic or transcriptomic studies in this species.

We unambiguously placed 13,470 *Alf*I fragments from the *de novo* reference in the genome assembly. We searched for and excluded the 15.2% of tags from the *de novo* reference that matched the reference matched more than one region equally well, and the 4.6% of genomic regions matched by more than one tag from the de novo reference. The 13,470 tags that remained include about half (52.8%) of the tags sequenced at sufficient depth for genotyping. We used these relationships between AlfI fragments in the de novo reference and genome assembly to assign each scaffold to a linkage group, capturing 304 scaffolds in this way. Because these scaffolds included some of the longest in the assembly (including 126 scaffolds > 1 Mb in length), this approach succeeded in capturing approximately two thirds of the total assembly (329 Mb, or 67.7% of the total). This resource substantially improves our understanding of long-range structural relationships in the *O. faveolata* genome. The average size of the 304 scaffolds integrated in the map was 1.05 Mb, and the average size of each collection of linked scaffolds was 17.1 Mb, a 16.3-fold increase in the length of chromosomal regions relative to analysis based on the assembly alone. The integrated map includes 8105 markers in 16 groups of linked scaffolds (506 markers per group on average), while the average scaffold in this resource includes only 30 markers (16.9-fold fewer). These comparisons serve to quantify the improvement in long-range structural information provided by the integrated map.

We assigned scaffolds from the genome assembly to the integrated map based on the structural framework established by the linkage map. We found 161 scaffolds containing a single marker, and assigned each to that marker’s position in the appropriate linkage group. Scaffolds containing more than one mapped marker presented an opportunity for evaluating agreement between these independent analyses. We found 143 scaffolds containing more than one marker. In most cases (88 scaffolds, or 61.5%) all markers mapped to a single linkage group, indicating independent agreement between the assembly and linkage analysis supporting the relative positions of these markers. In the cases where markers on a single scaffold were mapped to multiple linkage groups (n=55, 38.5% of the total), we assigned scaffolds to the group supported by most markers (placing 24 scaffolds), and leaving unplaced any scaffolds associated equally with more than one group (31 scaffolds). These conflicts between linkage analysis and genome assembly probably result in part from errors in each resource. Rather than attempting to resolve (in this study) all conflicts between these resources, our approach is to identify consensus regions supported by both analyses to develop a practical resource for genomic studies of *O. faveolata*.

Because the map only includes markers that were polymorphic and informative in the chosen cross, only a relatively small fraction of the markers genotyped in these samples (8%) were directly incorporated in the map. We drew on the relationships between markers, transcripts, and scaffolds to integrate additional markers into the map (Fig 1). We added markers mapping to the same scaffolds as markers already in the map (assigning these the same position in the map, since the distance between such closely linked markers cannot be accurately estimated from the number of offspring analyzed here). We added 7,107 markers to the map in this way, a 7.1-fold increase in the total number of markers in the integrated map.

Altogether, this process produced an integrated map containing 8,630 markers, 304 scaffolds, and 19,407 genes in a collection of 16 linkage groups. This resource is provided as Supplementary Table S1. In its present state the resource remains incomplete, since multiple genes and markers remain unplaced in the resource. The integrated map described here is not intended as a final version, but as an initial framework for genomic studies of *O. faveolata* that can easily be extended in future studies. Additional linkage maps can be used to identify relationships between markers and scaffolds, and subsequent improvements to the genomic assembly will naturally draw additional genes and markers into the integrated map.

### Genetic associations with variation in bleaching responses

To demonstrate the utility of the integrated map for genomic studies in *O. faveolata*, we conducted pilot studies of variation in bleaching responses during thermal stress experiments. Exposure to 4 degree heating weeks (DHW) of cumulative thermal stress induced substantial bleaching in coral fragments in the stress treatment, while corals in control conditions remained healthy and unbleached (Fig 3a). To analyze variation in bleaching responses of heat-stressed corals, we calculated bleaching scores for each fragment as the difference from the average color score of control fragments from the same colony. We partitioned variation in bleaching scores in a mixed model including origin and colony, revealing that bleaching responses varied significantly by origin (P=0.012) and by colony (P<0.001), explaining 47% and 33% of the variation in bleaching responses, respectively.

**Fig 3.**
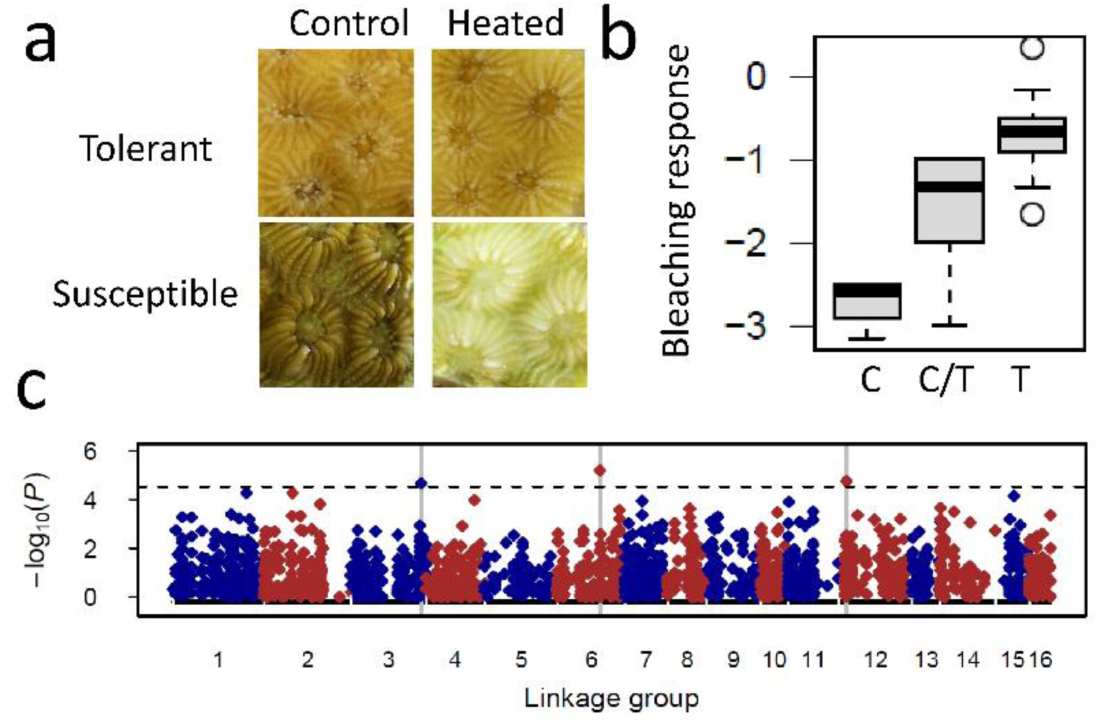
The integrated map can be used to study genomic association with variation in bleaching responses. This small-scale survey identified three markers associated with bleaching responses. (a) Examples of variation in bleaching responses observed during controlled thermal stress experiments, showing replicate fragments from susceptible and tolerant colonies after incubation in thermal stress or control treatments. (b) Effects of genotype on bleaching responses at marker on LG6. (c) Three markers (vertical grey lines) on LG 3, 6, and 12 were significantly associated with variation in bleaching responses during Thermal stress (FDR<0.05).

To control for confounding effects of variation in symbiont communities that may contribute to variation in thermal tolerance of the holobiont, we surveyed the dominant symbiont types in each colony by sequencing ITS2 amplicons using Sanger sequencing, and comparing the resulting sequences with ITS2 sequences from Symbiodinium clades A-D. Sequencing produced clean chromatograms for 16 of 24 samples, consistent with symbiont communities dominated by a single symbiont type. The remaining 8 samples consistently amplified well in PCR but produced chromatograms with overlapping peaks that prevented basecalling, as would be expected in mixed symbiont communities. On the basis of these sequence comparisons we annotated the dominant symbiont type in each sample as type B, D, or mixed. We observed clades B and D at intermediate frequencies across both populations of origin (clade B detected in 34% of FL samples and 56% of TX samples). Including this information in the mixed model revealed that variation in dominant symbiont types had no significant effects on bleaching responses in these samples (P=0.17).

#### Genetic markers associated with variation in bleaching responses

To understand the genetic basis for variation in bleaching responses among colonies, we conducted multilocus SNP genotyping of each colony, sequencing 2bRAD libraries to produce 2.8 million high-quality reads on average for each sample. We aligned these reads to the same reference developed from larval samples that naturally lack symbionts, to ensure that the loci genotyped were from the coral host rather than their algal symbionts. This analysis determined genotypes for >600 kb at ≥5× coverage. After filtering genotypes to minimize genotyping errors and missing data, we identified a set of 6,788 polymorphic markers (SNPs) genotyped in at least half of the samples.

To identify genomic regions associated with variation in bleaching responses, we tested for associations between bleaching responses and genotypes at each of these markers using mixed models controlling for genetic relationships among samples. For this analysis of genomic regions, we focused on the subset of 1,873 markers in the integrated map that were genotyped and polymorphic in these samples, mapping the statistical associations between bleaching and genotypes on to the integrated map (Fig 3). We found three markers (on LG 3, 6 and 12) were significantly associated with bleaching responses after multiple test correction, and were clearly elevated above the genomewide background (Fig 3).

#### Genes associated with variation in bleaching responses

Without the integrated map, the functional interpretation of these associations would be very limited, since none of the markers associated with bleaching responses directly matched gene sequences. Mapping these SNPs to the draft genome assembly made it possible to identify genes on the same scaffold as each SNP associated with bleaching. The number of genes identified in this analysis depended on the length of each scaffold. The number of genes identified in this analysis varied with scaffold lengths, finding 36-38 linked genes on the shorter scaffolds and 211 on the longer scaffold (Table 1). In contrast, analyzing the same data in the context of the integrated map allowed us to identify 5.6-fold more genes linked to each marker (range: 256-437 genes per marker) than the scaffold-based analysis (Table 1 & Supplementary Table S3).

**Table 1.** Analysis of association studies using the integrated map identifies more genes associated with phenotypic variation than sequence comparisons alone. Genes in each region are listed in Supplementary Table S3.

| Effects | Marker | LG | Scaffold | Genes directly matching markers | Genes on scaffolds with markers | Genes within 10 cM of markers |
| --- | --- | --- | --- | --- | --- | --- |
| Bleaching | denovoLocus12618 | 3 | MZGG01001646.1 | 0 | 36 | 290 |
|  | denovoLocus8166 | 6 | MZGG01001695.1 | 0 | 38 | 256 |
|  | denovoLocus9009 | 12 | MZGG01001743.1 | 0 | 211 | 437 |
| Fluorescence | denovoLocus9581 | 6 | MZGG01001262.1 | 1 | 118 | 118 |
|  | denovoLocus8443 | 6 | MZGG01000595.1 | 0 | 34 | 175 |
|  | denovoLocus1252 | 6 | MZGG01000597.1 | 0 | 67 | 256 |
|  | denovoLocus7916 | 10 | MZGG01001827.1 | 0 | 182 | 550 |
|  | denovoLocus12839 | 10 | MZGG01001871.1 | 0 | 61 | 235 |
|  | denovoLocus3937 | 11 | MZGG01001350.1 | 1 | 98 | 447 |
|  | denovoLocus10835 | 16 | MZGG01001189.1 | 1 | 59 | 511 |

Interestingly, each of these regions shared several functional categories of genes. Each region included multiple genes beta adrenergic receptor genes (3 on LG3, 2 on LG6, and 3 on LG12). Each region includes collagen genes (1 on LG3, 7 on LG6, and 7 on LG12). Each region also included multiple genes associated with ubiquitin-mediated protein degradation. This included five ubiquitin-protein ligase genes on LG3; three ubiquitin-protein ligase gene and one ubiquitin hydrolase gene on LG6; and three ubiquitin-protein ligase genes and six ubiquitin hydrolase genes on LG12 (Supplementary Table S3). We also found multiple genes with known roles in thermal stress responses, including eight genes on LG3 homologous to universal stress proteins and one homologous to Hsp70. This analysis of genes linked to the bleaching associated marker greatly increases the number of functional hypotheses generated from this dataset relative to an analysis based only on the genome assembly, illustrating the value of the integrated map developed here for studying genomic process and patterns in *O. faveolata*.

### Genetic associations with fluorescence phenotypes

The coral colonies chosen for our pilot study also presented conspicuous variation in color, resulting from differences in the intensity of green fluorescence from endogenous fluorescent proteins (GFPs) (Fig 4a). Since this additional trait presents another opportunity to study the genomic basis for variation in the trait, we quantified this variation using fluorescence microscopy, and partitioned variation in fluorescence into components of colony and origin. This analysis revealed significant variation among colonies (p<0.001), explaining 80.3% of variation, while origin (FL or TX) had no effects on fluorescence.

**Fig 4.**
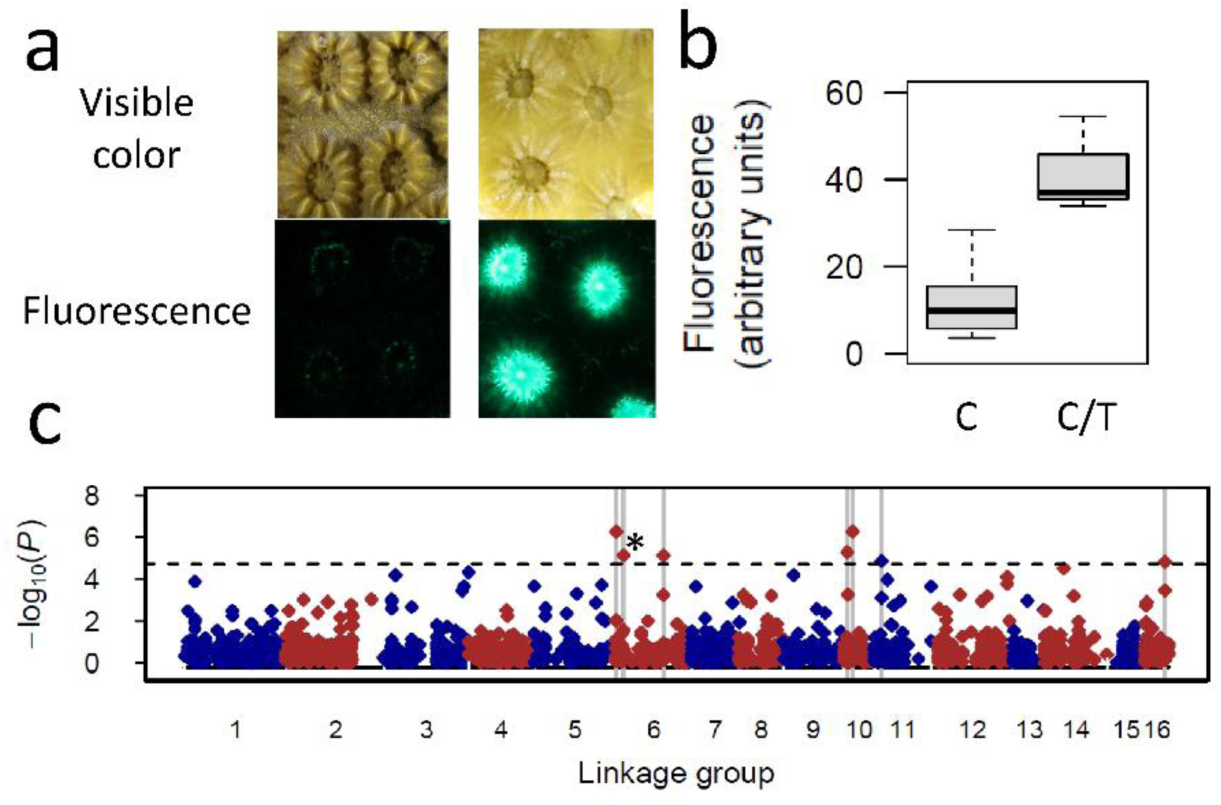
The integrated map can be used to study genomic associations with variation in color phenotypes. Here we identified four markers significantly associated with variation in green fluorescence. (a) Examples of color variation resulting from variation in fluorescent proteins. Fragments with contrasting bright and dim phenotypes are shown under white light (top), and under fluorescence microscopy (excitation = 470 nm) (bottom). (b) Effects of genotype at a candidate SNP on LG 6 (* in panel c) on fluorescence. (c) Genomic analysis identifies seven markers (vertical grey lines) associated with fluorescence intensity (FDR<0.05).

#### Genomic regions associated with color variation

To identify genomic markers and regions associated with this variation, we tested for associations between fluorescence and genotype at each marker using a series of linear models, as described above for analysis of bleaching responses. We focused on the same subset of 1,877 markers in the integrated map that were genotyped in these samples, mapping the statistical associations between fluorescence and genotypes onto the integrated map (Fig 4b,c). This analysis identified regions on LG 6, 10, 11, and 16 associated with fluorescence.

#### Genes associated with color variation

Few of the markers associated with fluorescence matched transcripts directly, obscuring functional implications of these associations (Table 1). Expanding our view to consider the entire scaffold containing each gene, we identified genes linked to each marker with the number of genes correlated to the length of each scaffold (34-67 genes on shorter scaffolds, and 98-182 on longer scaffolds). Comparison with the integrated map expanded information on the genes linked to each marker substantially, identifying 118-550 genes for each marker (Table 1 & Supplementary Table S4).

Interestingly, none of these genes match fluorescent proteins themselves. The integrated map includes 13 genes annotated as fluorescent proteins (on LG1 and 4), but none are located near markers that were significantly associated with variation in fluorescence of our samples. Instead, we found multiple regulatory genes (Supplementary Table S4). These results suggest that variation in regulatory genes rather than the fluorescent proteins themselves contributes to the variation in fluorescence intensity among our samples. We evaluated word frequencies in functional annotation of genes in these regions and found that “receptor” was the most frequent informative term (138 genes). While the breadth of this category limits functional inferences from this observation, the group included several specific receptors that were repeatedly linked to fluorescence-associated markers on multiple linkage groups. We found galanin receptor genes linked to fluorescence-associated markers (FAMs) on LG6, 10, 11, and 16. We also found histamine receptors linked to FAMs on LG6, 10, and 16, and neuropeptide receptors linked to FAMs on LG 10, 11, and 16. The repeated observation of genetic variation linked to these receptors suggests the possibility that variation in these receptors contributes to variation in fluorescence, possibly by mediating corals’ regulatory responses to environmental and cellular factors promoting expression of fluorescence proteins.

Another striking pattern apparent from word frequencies was the abundance of growth factor and growth factor receptor genes linked to FAMs on multiple linkage groups. We found multiple growth factor receptor genes linked to FAMs on each of LG 6 and 10, and multiple growth factor genes on each of LGs 10, 11, and 16. These patterns suggest the possibility that variation in growth factor-mediated signaling contributes to variation in fluorescence in our samples. The novel patterns identified in this study provide new hypotheses for the striking variation in color morphs observed in natural populations, and illustrates the utility of the integrated map for genomic studies of trait variation in *O. faveolata*.

### Genomic analysis of population divergence

Finally, we analyzed variation in the same set of SNP genotypes among a collection of corals originating from distant populations in the Florida Keys and the Flower Garden Banks (Gulf of Mexico). Considering the entire set of 2,905 markers that were integrated into the map and genotyped at sufficient coverage in these samples for estimates of F_ST_, we found substantial divergence between these groups (average F_ST_ = 0.162). Locus-specific estimates showed a broad distribution of F_ST_ values, with little divergence at most loci (e.g. 57% of markers had F_ST_ < 0.1, and 81% of markers showed F_ST_ < 0.3). The small number of outlier loci with high F_ST_ may result from differences in selective pressures or demographic histories of these populations, and mapping these markers onto the integrated map allowed us to investigate the relative positions of these highly-differentiated markers in the genome. We used a sliding window approach to investigate these patterns, revealing that the most strongly differentiated markers are clustered in a few genomic regions. Estimating F_ST_ in 25-marker windows across the map, we found that F_ST_ estimates for most regions were near the genome-wide average of 0.16, while the most strongly differentiated 1% of windows were clustered in five regions on LG 12, 13, 14, and 16 (Fig 5). These regions showed extremely high differentiation, with average F_ST_ values exceeding 0.31.

**Fig 5.**
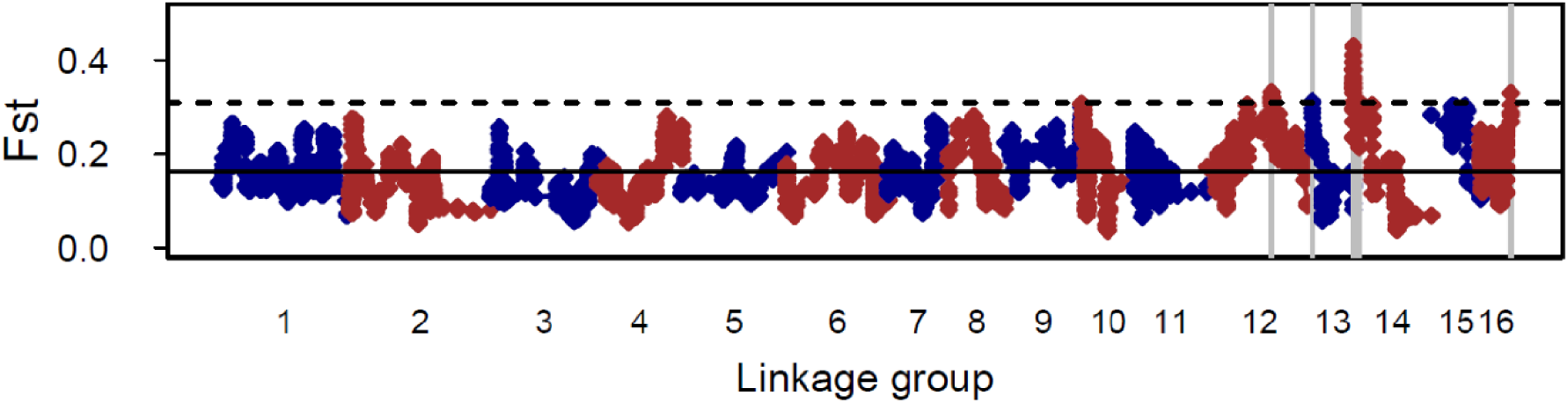
The integrated map can be used to study genomic patterns of genetic differentiation between populations. This sliding window analysis of corals from FL and TX identified a few genomic regions that were more strongly differentiated between populations than the overall genomic background. Each symbol depicts the average F_ST_ calculated for a window of 25 markers. The solid horizontal line depicts the genomewide average F_ST_ (0.162), and the dashed line shows the threshold (F_ST_ = 0.31) delineating the most highly differentiated 1% of genomic regions (highlighted with grey vertical lines).

We used the integrated map to investigate gene content of these highly differentiated regions. Altogether these regions included 466 annotated genes, providing a rich resource for investigating the functional context of genetic divergence between populations (all genes shown in Supplementary File S5). We found multiple zinc finger metalloproteinase genes in differentiated regions on both LG 12 and 14. We found multiple neuronal acetylcholinesterase receptor genes in differentiated regions on LG 12 and 14, and numerous transcription factors in differentiated regions on LG 12, 13, and 14. Our findings demonstrate that the most strongly differentiated regions of the genome include multiple regulatory genes. Determining the adaptive significance of this variation is outside the scope of this study, but our analysis demonstrates the benefits of the integrated map for population genomic studies in *O. faveolata*. On the own, anonymous SNP markers would be limited to descriptions of average genomewide differentiation among populations, while genomic analysis using the integrated map provides insights into the possible functional consequences of that differentiation.

## DISCUSSION

We developed a genetic linkage map for the coral *O. faveolata* as a resource for studying genomic patterns of trait association or genetic differentiation in this system. Here we have described the linkage map and our efforts to integrate this resource with existing sequence assemblies, and demonstrated the utility of these resources for studying trait associations and genetic differentiation in these corals. This allowed us to expand our analysis beyond direct sequence comparisons of genetic markers, transcripts, and scaffolds to include long-range structural information and identify a far greater number of genes associated with this variation. Future genomic studies of variation in natural populations of *O. faveolata* will benefit from mapping statistical associations onto this same resource, providing a common framework for comparing signals from different studies, traits, or populations to identify genes and markers reproducibly associated with this variation.

While the haploid number of chromosomes in *O. faveolata* has not been empirically determined, our analysis of recombination frequencies identified 16 linkage groups that we hypothesize represent chromosomes or portions of chromosomes. For comparison, data on haploid chromosome numbers (karyotype) are available for multiple (31) coral species [20,69-72], and these range from n=12 to n=27 across all Scleractinian corals measured. The Scleractinian phylogeny includes two major clades (called robust and complex) that probably diverged >240 mya [73]. Only two of these species (*Fungia scutaria* and *Favia pallida*) are members of the robust clade like our focal species (*O. faveolata*), and both have haploid complements of 14 chromosomes. These comparisons place our estimate of 16 LGs well within the range of chromosome numbers for Scleractinians, although slightly higher than the 14 chromosomes reported in other robust corals.

Few direct comparisons are available for our linkage map, since only two other genetic linkage maps have been reported in corals, both in the widely studied *Acropora millepora* [31,32]. Our map includes 998 markers, compared with 429 in the map based on individually genotyped markers [32] and 1,458 in the map developed using 2bRAD [31]. Our total map length (2,049 cM) was substantially longer than previous studies (1,493 and 1,358 cM respectively). This difference may result from differences in mapping software (Onemap versus JoinMap) or genotyping errors in one or both datasets. Marker densities were comparable in all studies: 1 per 2.05 cM (present study), 1 per 3.4 cM [32], and 1 per 0.91 cM [31]. Both studies of *A. millepora* identified 14 linkage groups, consistent with expectations from karyotyping [72], and our analysis identified a comparable number of linkage groups (16) in *O. faveolata*.

In this study we developed an integrated map by combining our genetic linkage map with a recently released draft genome assembly. Similar resources are rapidly emerging for a wide variety of non-model species of interest for ecological or evolutionary studies. Alone, each of these resource types has limitations. Annotated transcriptome assemblies are widely available for corals and other non-model species [11,14-17,74,75]. However, while these resources are essential tools for many studies they offer no information on the relative positions of these genes in the genome. Draft genome assemblies are more expensive and computationally challenging than transcriptome assemblies, but these resources have also been rapidly emerging in recent years in many non-model systems including corals [19,20]. Initial draft assemblies produced from high-throughput DNA sequencing are typically fragmented, consisting of thousands of fragmented scaffolds with average lengths in the 10s of kb, rather than the 10s of Mb-scale scaffolds expected in finished assemblies. This limits the long-range structural information available in these resources, obscuring the relative positions of genetic markers across the genome. Finishing genome assemblies remains challenging, especially in highly polymorphic or repetitive genomes, and this challenge is the focus of extensive ongoing research using long sequencing reads or novel assembly algorithms to improve genome assembly, as recently reviewed in [76].

Here, we’ve used a comparatively older approach (genetic linkage mapping) to develop a structural framework for genomic studies in *O. faveolata*. Our approach was inspired by previous studies using high-throughput DNA sequencing to combine genomic assemblies and genetic linkage maps [77-79]. Relative to more tractable marine invertebrates, the restricted distribution and life history of reef-building corals have limited the development of genetic linkage maps in these systems. Although the reproduction of corals cannot be easily manipulated, the seasonality of annual mass-spawning events [80,81] does allow for accurate prediction of reproduction in many species, making it possible to conduct controlled genetic crosses using gametes from known parental colonies. Similar crosses have been used to study many aspects of coral biology including symbiosis [82], species boundaries [49,83], and responses to thermal stress [84-86], and we’ve extended this approach to measure quantitative genetic parameters and develop genetic linkage maps in corals [31,32,47]. The long generation times of most corals preclude multi-generation designs (F2 and beyond), so we used a mapping approach developed for linkage analysis in outcrossing plant species, which allows for linkage analysis in first-generation (F1) families produced by crossing outbred parents from natural populations [23]. Corals undergo extensive mortality during larval and post-settlement periods [87-89], which are likely to distort segregation ratios extensively in the survivors, like other marine invertebrate larvae [89,90]. To minimize these distortions, we focused on early developmental stages. These small larval stages (∼100 μm) contain little genomic DNA (∼200 ng larva^-1^), presenting technical challenges for preparation of sequencing libraries. To address this challenge we prepared sequencing libraries directly from alcohol-precipitated lysates of individual larvae, maximizing yields while still producing high-quality sequencing libraries. In subsequent experiments with other Cnidarian species we have found this procedure similarly effective even at lower DNA amounts (<100 ng).

Alone, the genetic linkage map provides little functional information for genomic studies. Few of the markers included in the map directly coding regions, so the map provides very little direct information on the genes associated with each genomic region. However, integrating linkage map and annotated genome assemblies made it possible to anchor 59.5% of the genes and 67.7% of the genome assembly in a long-range framework established from the genetic linkage map. While this initial version of the integrated resource remains incomplete, it has already substantially expanded the set of associations between genetic markers and genes available for genomic studies. Our demonstration studies of trait associations and population differentiation illustrate this benefit. Few of the markers associated with bleaching or fluorescence matched transcripts directly. However, mapping these results onto the integrated map identified hundreds of genes linked to each trait-associated marker. Similarly, few markers in the most highly-differentiated 1% of the genome matched genes directly, offering little information on the functional context for these patterns. In contrast, the integrated map revealed hundreds of genes in each highly-differentiated region. Genomic scans for signatures of differentiation or trait associations are powerful tools for hypothesis generation. However, in the absence of an integrated map, reduced representation approaches for SNP genotyping provide little functional information. Whole genome sequencing overcomes this limitation, but remains prohibitively expensive for widespread use as a genotyping tool in organisms with large genomes. Analysis of the transcriptome provides genetic information directly in functional coding sequences, but is limited by expression levels and complicated by allele specific expression. The approach we have outlined here, anchoring genes and genetic markers in an integrated resource developed from draft genome and transcriptome assemblies and a genetic linkage map, provides a framework for functional interpretation of genetic data from the widely used and cost-effective family of RAD and associated methods in non-model species.

## Acknowledgements

The authors are grateful to Monica Medina and Bishoy Hannah (Penn State) for sharing their draft genome assembly prior to publication. We thank Mikhail Matz (Univ. Texas – Austin), Sarah Davies (Boston University), and Carly Kenkel (Univ. Southern California) for assistance with sample collection and fieldwork logistics. We thank the staff of the Flower Garden Banks National Marine Reserve for supporting fieldwork and sampling on board the R/V Manta (Emma Hickerson, Marissa Nuttal, GP Schmahl). We thank Rebecca Vega Thurber for sharing coral specimens for thermal stress experiments.

## Authors’ contributions

JS prepared sequencing libraries for SNP genotyping. SG and KD conducted thermal stress experiments and helped with sample collection. EM conceived and supervised the study, conducted all linkage and genomic analysis, assisted with thermal stress experiments and sample collections, and wrote the manuscript. All authors read and approved the final manuscript.

